# Targeted preoptic neuronal activation induces a hypothermic state with metabolic reprogramming and acute cerebroprotection

**DOI:** 10.1101/2025.10.24.684192

**Authors:** Aizad Kamal, Juan Liu, Ernesto R. Gonzales, Khairunisa Mohamad Ibrahim, Fan Zhang, Tamara Abou-Antoun, Hannah E. Skelton, Javier Kelly Cuenca, Carla Yuede, Gary J. Patti, Leah P. Shriver, Jin-Moo Lee, Aaron J. Norris, Eric C. Landsness

## Abstract

**Objective:** Therapeutic hypothermia is cerebroprotective in experimental stroke but has shown limited clinical efficacy due to metabolically costly counter regulatory responses. Naturally occurring states such as torpor achieve hypothermia through coordinated regulation of temperature and metabolism and are associated with cerebroprotection. Here, we tested whether selective activation of a genetically defined thermoregulatory neuronal population is sufficient to induce hypothermia after ischemic injury and whether temperature-dependent cerebroprotection is accompanied by metabolic remodeling. Using chemogenetic tools, kappa opioid receptor expressing neurons in the preoptic area were activated in mice subjected to transient middle cerebral artery occlusion. Core body temperature, metabolism, infarct volume, and motor recovery were assessed under conditions permitting hypothermia or maintaining normothermia. Untargeted metabolomic profiling was performed on whole brain tissue collected 30 minutes after activation in the absence of ischemia. Activation of kappa opioid receptor expressing preoptic neurons induced a hypothermic state with coordinated metabolic changes, reduced infarct volume and improved acute motor outcomes. These effects were attenuated when body temperature was maintained at normothermia. Metabolomic profiling revealed coordinated alterations in nucleotide, phospholipid, and sphingolipid pathways. These findings link temperature dependent cerebroprotection to coordinated metabolic remodeling, with translational relevance for stroke.

## Introduction

Therapeutic hypothermia improves neurological outcomes and survival in patients with cardiac arrest^1,2^, in neonates with hypoxic-ischemic encephalopathy^3^ and confers robust cerebroprotection in preclinical models of stroke^4–7^. Despite this consistent biological efficacy, therapeutic hypothermia has not demonstrated benefit in clinical trials of ischemic stroke^8–11^. In endotherms, including mice and humans, imposed cooling activates homeostatic defenses that counter reductions in body temperature^12^. These defenses are metabolically demanding, requiring substantial energy expenditure^12,13^, and engage systemic metabolic pathways that may interfere with—or obscure—the processes underlying hypothermic protection. Together, these observations suggest that hypothermic efficacy may not depend on temperature reduction alone, but on engagement of coordinated physiological and metabolic programs that shape tissue responses during ischemia.

Consistent with this concept, naturally occurring hypothermic states such as torpor and hibernation coordinate suppression of body temperature, metabolism, and energy expenditure^14^, and confer robust cerebroprotection in preclinical models^15,16^. Recent brain-tissue metabolomic and lipidomic studies of natural torpor and hibernation reveal coordinated remodeling of metabolic pathways, supporting the view that torpor represents a distinct, actively regulated metabolic state^17,18^. However, it remains unclear whether the coordinated metabolic remodeling in hypothermic states reflects reduced body temperature itself or the neural programs that initiate and maintain it. Defining the metabolic organization of hypothermic states and identifying the neuronal populations that coordinate temperature and metabolism are therefore essential to understanding their contribution to cerebroprotection.

The preoptic area (POA) contains functionally distinct neuronal populations linked to multiple homeostatic processes, including regulation of body temperature and organism-level metabolism in response to environmental temperature and energy availability, positioning the POA as a central regulator of thermoregulatory physiology^19,20^. In mouse models of stroke, stimulation of the POA induces hypothermia, suppresses counter-regulatory responses and robustly reduces ischemic brain injury^21,22^. However, these studies have largely relied on broad engagement of POA neuronal ensembles, leaving unresolved whether activation of specific, genetically defined neuronal subpopulations within the POA is sufficient to induce a hypothermic and hypometabolic state. Recent work, including anatomical, physiological, and systems-level characterization from our group and others, has identified a genetically identifiable subpopulation of POA neurons expressing the kappa opioid receptor (KOR) that plays a central role in regulating thermoregulatory and metabolic state. Activation of these KOR-expressing neurons (POA^KOR^⁺) induces a profound hypothermic and hypometabolic state, demonstrating that selective engagement of a discrete subpopulation is sufficient to suppress body temperature and regulate metabolism^23,24^. Importantly, KOR-expressing neurons have also been identified within the human POA, suggesting that key elements of this thermoregulatory architecture are conserved across species^25,26^. This body of evidence positions POA^KOR^⁺ neurons as a genetically defined, evolutionarily conserved, and experimentally targetable neuronal subpopulation for probing coordinated hypothermia and defining its metabolic architecture.

In this study, we tested whether selective activation of POA^KOR^⁺ neurons, is sufficient to induce a metabolically coordinated hypothermic state that is associated with acute cerebroprotection. Using chemogenetic activation in a model of transient cerebral ischemia, we assessed physiological and functional outcomes under conditions permitting or preventing hypothermia to examine the contribution of temperature to cerebroprotection. In parallel, we performed untargeted metabolomic profiling to characterize the biochemical organization of the induced hypothermic state. We found that activation of POA^KOR^⁺ neurons produced a coordinated hypothermic, metabolically reprogrammed state; cerebroprotection occurred only when hypothermia was permitted; and this state was accompanied by temperature-dependent remodeling of nucleotide, phospholipid, and sphingolipid pathways.

## Methods

### Overall Experimental Strategy

This study consisted of three complementary experiments designed to determine whether selective activation of a genetically defined thermoregulatory neuronal population induces a coordinated hypothermic state associated with metabolic remodeling and acute cerebroprotection after ischemic injury. Across all experiments, KOR-Cre mice first underwent stereotaxic viral delivery targeting POA^KOR+^ neurons and were allowed three weeks for DREADD expression. At three weeks after viral injection, body temperature monitoring devices were implanted.

Following a one-week recovery period, mice received a single validation dose of clozapine-N-oxide (CNO; 0.5 mg/kg) to confirm successful chemogenetic activation of POA^KOR+^ neurons based on reductions in body temperature and behavioral activity. Mice were then allowed an additional one-week recovery period before being randomized to the experimental protocols described below. Each animal received a single experimental CNO injection (0.5 mg/kg) according to the respective study design. Repeated CNO injections were not used during the experimental protocols. Across all experiments, ambient temperature was manipulated to either permit or suppress the fall in core body temperature, allowing assessment of the contribution of reduced body temperature independent of continued POA^KOR^⁺ neuronal activation.

<u>Experiment</u> 1 examined whether the coordinated hypothermic and hypometabolic response to POA^KOR+^ activation, previously characterized in non-ischemic mice^24^, remained present following ischemic stroke. Mice underwent transient middle cerebral artery occlusion (tMCAO) followed by either a control condition or POA^KOR+^ activation under conditions permitting hypothermia. Twenty-four hours later, serum and liver tissue were collected for targeted metabolomic analysis focused on glycolysis, ketone metabolism, tricarboxylic acid (TCA) cycle intermediates, and energetic metabolites.

<u>Experiment</u> 2 examined whether the hypothermic state induced by POA^KOR+^ activation was associated with acute cerebroprotection and whether these effects were temperature dependent. Mice underwent tMCAO followed by assignment to one of three conditions: (1) control, (2) hypothermic-POA^KOR+^ activation under thermoneutral housing conditions (30 °C), or (3) normothermic-POA^KOR+^ activation under elevated ambient temperature (36 °C) to suppress hypothermia despite continued neuronal activation. Core body temperature was continuously monitored for 72 hours, followed by behavioral testing and infarct quantification.

<u>Experiment 3</u> examined metabolic remodeling in the brain 30 mins after activation of POA^KOR+^ neurons in the absence of ischemic brain injury. Mice expressing Gq DREADDs or eYFP in POA^KOR+^ neurons received CNO injections and whole brains were collected 30 minutes later for untargeted metabolomic and lipidomic profiling to characterize temperature-associated metabolic organization. This experiment specifically focused on brain metabolic remodeling associated with the cerebroprotective state, and metabolomic profiling was therefore restricted to brain tissue. This design isolated early metabolic effects of POA^KOR⁺^ activation and hypothermia prior to the confounding effects of ischemic injury, inflammation, or tissue necrosis.

Shared experimental procedures for viral targeting, temperature monitoring, ischemic stroke induction, behavioral assessment, metabolomic acquisition, and statistical analysis are described below.

### Mice

Adult (9–10 weeks old at start of experiments) KOR-Cre strain mice (RRID:IMSR_JAX:035045) were used in all experiments. Prior to the experiment, all mice were kept on a 12-hour light/12-hour dark cycle, with a room temperature of 22° ± 1 °C with food and water available to them ad libitum. Animals were randomly assigned to experimental groups following confirmation of viral expression and prior to experimental procedures. Both male and female mice were included in all experiments and sex was not used as an exclusion criterion.

Investigators remained blinded to group allocation during data collection and analysis. Throughout the ensuing testing and analysis, experimenters were blinded to the treatments given to the mice and the processing of the samples. Mice were monitored daily with ad libitum access to food and water. Animals were monitored daily for postoperative recovery and overall health status, including body weight, hydration, grooming, mobility, and neurological condition.

Humane endpoints were determined according to institutional animal care and guidelines, and animals showing signs of severe distress or compromised welfare were euthanized.

### Animal Ethics Statement

All animal procedures were approved by the Washington University School of Medicine Institutional Animal Care and Use Committee (IACUC). All experiments were conducted in accordance with the Guide for the Care and Use of Laboratory Animals (8th edition, National Research Council, National Academies Press, 2011) and were reported in compliance with ARRIVE 2.0 (Animal Research: Reporting of In Vivo Experiments) guidelines.

### Chemogenetic Viral expression

At 9-10 weeks of age, experimental mice were bilaterally injected with the Gq virus, AAV5/hSyn-DIO-hm3D(Gq)-mCherry (7.8 × 10¹² vg/ml, Addgene) + AAV5-EF1a-DIO-hChR2(H134R)-EYFP (2.5 × 10¹³ vg/ml) with a 1:1 ratio, using Neuros Syringe (65457-01, Hamilton Com.) with a volume of 100 nl on both sides of the POA (AP +0.40 mm, ML ±0.45 mm, DV-5.35 mm) with an injection speed of 30 nl/minutes. Control mice were injected with AAV5-EF1a-DIO-eYFP (1.8 × 10¹³ vg/ml), with the same volume, brain coordinates and injection speed as the experimental group. Four weeks following viral injection, after a one-week recovery period following implantation of the body temperature monitoring device, a single test dose of clozapine-N-oxide (CNO; 0.5 mg/kg) was administered to confirm functional engagement of POA^KOR+^ neurons, as evidenced by a reduction in body temperature and behavioral activity. Animals that did not achieve a ≥3 °C reduction in core body temperature within 30 minutes following CNO administration were excluded based on pre-specified validity criteria.

### Body Temperature Monitoring

Three weeks following viral injection, mice were sedated with 1.5% isoflurane before implanting an iButton data temperature logger (Thermocron DS1925L#50, running OneWireViewer software, version 3.19). The skin was cut with a minor incision, and the iButton was placed at the dorsal part of the body lateral to the midline after the dermal space was opened. The mice’s skin was sutured together after the surgery using 1-2 wound surgical clips, that were then removed one week later. Alternatively, mice were implanted with wireless IPTT-300 temperature transponders (Bio Medic Data Systems; Seaford, DE) while anesthetized with 2% isoflurane. The transponder was implanted beneath the skin, rostral to the right hind leg. This approach for continuous temperature monitoring using implantable transponders and iButton data loggers has been validated as a reliable method for measuring body temperature in small animal models, with these devices demonstrating strong correlation with core temperature measurements and high accuracy under physiologically relevant conditions^27,28^. In addition, in a separate validation cohort, brain temperature was measured using a fine-wire K-type thermocouple probe (0.1-mm diameter) while body temperature was simultaneously monitored using the implanted temperature logger. Simultaneous measurements of brain temperature and body temperature obtained demonstrated closely parallel temperature declines following a single CNO administration (Supplementary Fig. S2).

### Ischemic stroke

Mice underwent tMCAO, as previously reported^29^. Under 2% isoflurane, the left middle cerebral artery (MCA) was exposed by a temporal muscle incision, and baseline blood flow in the cortical MCA region was measured using laser Doppler flowmetry (LDF; Model 403A, Transonic Systems, Inc TSI, Inc.). A ventral midline incision was made in the neck to expose the left common carotid artery. To temporarily block the origin of the MCA, a silicon-coated, 6.0-gauge, nylon suture, measuring 12 mm in length, was inserted into the carotid artery approximately 9.0–10.5 mm in depth. A thermoregulated heating pad was used to maintain normal body temperature of 36 °C during the surgical process. To restore cerebral circulation, the suture was removed after 30 minutes. Cerebral perfusion (% baseline) was monitored pre-occlusion, every 10 minutes during occlusion, and within 5 min of reperfusion and the laser Doppler probe was removed at the completion of surgery. A separate perfusion measurement was performed at the 72-hour endpoint by repositioning the probe over the same cortical region using the original stereotaxic landmarks (Supplementary Fig. S1).

Animals were excluded based on pre-specified validity criteria related to adequacy of ischemia and reperfusion. These criteria included failure to reach ≤15% of baseline perfusion during tMCAO, failure to recover ≥50% of baseline flow within 5 minutes of reperfusion, or failure to recover ≥70% of baseline flow at 72 hours.

### Behavior

#### Rotarod

Mice were placed on a rotarod set to a constant speed of 4 rotations per minute for 5 minutes for a single trial. The time it took for the mouse to drop off the drum was recorded (maximum 300 seconds). Testing was performed before (baseline) and 72 hours after tMCAO (post 72 hours) when animals were available for behavioral assessment.

#### Pole

Mice were positioned with their head pointing up at the top of a 60 cm vertical pole with a 1 cm diameter. As soon as the mouse started to turn, the recording started. The total time (Ttotal) to drop to the floor and the time (Tturn) to turn entirely to face down the pole were recorded. A maximum score was given when a mouse instantly fell off the pole. Each mouse was tested for two trials in total. The average scores of trial one and trial two were used for data analysis. Testing was performed before (baseline) and 72 hours after tMCAO (post 72 hours) when animals were available for behavioral assessment.

#### Corner

Two pieces of cardboard (30 cm × 20 cm × 1 cm) were placed at a 30-degree angle to create a corner. Then the mouse was placed 1 cm away from facing the corner. A trial was recorded when the mouse moved towards the corner, and it reared in response to whisker stimulation before turning back towards the open end. The direction it turned was then recorded as left or right.

Each mouse had a total of 10 trials recorded and an asymmetry score was calculated (Right turn/(Right turn+Left turn) *100). Testing was performed before (baseline) and 72 hours after tMCAO (post 72 hours) when animals were available for behavioral assessment.

### Stroke Size

Seventy-two hours after tMCAO each mouse was euthanized under deep anesthesia using a full dosage of 5% isoflurane. Mice were then given an intraperitoneal euthanizing dosage of sodium pentobarbital at a concentration of 100 mg/mL per-mouse. Perfusion and isolation of the heart was done accordingly. After euthanasia and intracardiac perfusion, mice brains were removed from the skull sliced into 1 mm-thick sections, and subsequently stained in 2,3,5-triphenyltetrazolium chloride (TTC) at 37 °C for 5 minutes. The primary endpoint was acute infarct volume at 72 hours following transient MCAO. Accordingly, TTC staining was selected because it remains the field-preferred histological method for quantifying infarct volume during this early post-ischemic period, with extensive validation demonstrating excellent agreement with conventional histological assessments at this time point^30,31^. After TTC staining, each brain sliced images were digitized and analyzed using Adobe Photoshop, and then stored in 4% paraformaldehyde. Direct infarct volume within the ipsilateral hemisphere was quantified using SigmaPlot 11.2 (Systat Software, San Jose, CA, USA) using a similar previously established protocol^29^ for animals in which brain tissue was available for TTC analysis. One animal in the hypothermic-POA^KOR^⁺ group died prior to the planned 72-hour endpoint. However, brain tissue was recovered immediately after death and included in the TTC infarct volume analysis. Therefore, the TTC analysis included 14 animals despite 3 mortality events being recorded in this group.

### Serum and Liver Metabolite Analysis

All tissues were collected using an identical standardized protocol and processed identically across experimental groups to minimize pre-analytical variability. Blood was collected at the time of euthanasia and serum isolated by centrifugation. Liver tissue was rapidly harvested and flash-frozen in liquid nitrogen prior to storage at −80 °C. Polar metabolites were extracted from liver tissue using a 2:2:1 mixture of acetonitrile:methanol:water as previously described^32,33^.

Briefly, liver tissue was weighed and homogenized in extraction solvent using ceramic beads and a bead mill homogenizer (Omni International). Samples were incubated at −20 °C for two hours followed by centrifugation at 14,000 revolutions per minute (RPM) for 10 minutes at 4 °C. Supernatants were transferred to autosampler vials and stored at −80 °C until analysis Polar metabolites and lipids were extracted from serum using solid-phase extraction (SPE). An aliquot of serum was pipetted into the well of the SPE plate (Agilent Captiva EMR-Lipid, 1 mL 96-well plate, 40 mg sorbent mass, Agilent Technologies, Santa Clara, CA) mounted over a 1 mL collection plate. A solution consisting of ice-cold 1:1 acetonitrile:methanol was added to the wells in a 1:4 ratio and the plate was shaken for 1 minute at room temperature at 360 RPM on an orbital shaker. The plate was then incubated at 4° C for 10 minutes. Extraction solution consisting of 2:2:1 acetonitrile:methanol:water was added to the wells in a ratio of 1:5 serum to extraction solution. The plate was shaken for 10 minutes at room temperature at 360 RPM on an orbital shaker. A second volume of extraction solution was added to the plate in a ratio of 1:1 serum to extraction solution and eluted from the plate by using a positive pressure manifold (PPM-96, Agilent Technologies, Santa Clara, CA). The collection plate was sealed with foil (AlumaSeal II Aluminum Adhesive Sealing Foil, Research Products International Corp, Mount Prospect, IL) and stored at −80 °C until analysis. The lipid fraction of the serum was eluted from the SPE plate into a new 1 mL collection plate by two additions of 500 uL of 1:1 mix of methyl tert-butyl ether and methanol to the plate followed by elution with the positive pressure manifold. The lipid fraction was dried-down under nitrogen (Biotage SPE Dry 96 Dual, Biotage, Uppsala, Sweden) and the plate was sealed with foil and stored at −80 °C until analysis. To reconstitute for analysis, a reconstitution solution of 1:1 methanol:isopropanol was added to the collection plate in a ratio of 1:4 serum to reconstitution volume. Ultra-high performance liquid chromatography coupled with mass spectrometry (UHPLC/MS) analyses were performed using a Thermo Scientific Vanquish Flex UHPLC system interfaced with a Thermo Scientific Orbitrap ID-X Mass Spectrometer. Polar metabolites were separated using a HILICON iHILIC-(P) Classic HILIC column (100 × 2.1 mm, 5 µm) with a corresponding guard column (20 × 2.1 mm, 5 µm). Lipids were separated by using an ACQUITY Premier HSS T3 column (100 x 2.1mm, 1.8µm, Waters). The mobile-phase solvents consisted of solvent A = 20 mM ammonium bicarbonate, 2.5 µM medronic acid, 0.1% ammonium hydroxide in 95:5 water:acetonitrile and solvent B = 2.5 µM medronic acid in 95:5 acetonitrile:water. The column compartment temperature was maintained at 45 °C, and metabolites were eluted using a linear gradient at a flow rate of 250 µL/ minute. Following LC/MS analysis, a targeted list of metabolites glycolysis, ketogenesis, TCA cycle metabolism, and cellular energy balance were analyzed by using Skyline^34^.

### Untargeted Brain Metabolomics and Lipidomics

All tissues were collected using an identical standardized protocol and processed identically across experimental groups to minimize pre-analytical variability. Following live decapitation, whole brains were immediately removed from the skull, flash-frozen in liquid nitrogen to halt metabolism, and stored at −80 °C until polar and lipid metabolite extraction as previously described^32,33^. Briefly, the brain tissue was weighed and added to a 15 mL falcon tube containing 2.8 mm ceramic beads. A 2:2:1 mixture of acetonitrile:methanol:water (40 µL per 1 mg of tissue) was added and the samples were homogenized using the tissue-specific presets on a bead mill (Omni International). Samples were incubated at −20 ° C for one hour followed by centrifugation at 10,000 RPM for 10 minutes at 4° C. The extract was transferred to autosampler vials and stored at −80° C until analysis. UHPLC/MS analyses were conducted by using a Thermo Scientific Vanquish Flex UHPLC system, interfaced with a Thermo Scientific Orbitrap ID-X Mass Spectrometer. A HILICON iHILIC-(P) Classic HILIC column (100 x 2.1 mm, 5 µm) with a HILICON iHILIC-(P) Classic guard column (20 x 2.1 mm, 5 µm) was utilized for the separation of polar metabolites. The mobile-phase solvents consisted of solvent A = 20 mM ammonium bicarbonate, 2.5 µM medronic acid, 0.1% ammonium hydroxide in 95:5 water:acetonitrile and solvent B = 2.5 µM medronic acid in 95:5 acetonitrile:water. The column compartment temperature was maintained at 45 °C, and metabolites were eluted using a linear gradient at a flow rate of 250 µL/ minute as follows: 0-1 min, 90% B; 12 minutes, 35% B; 12.5-14.5 minutes, 25% B; 15 minutes, back to 90% B. For the separation of lipids, an Acquity UPLC HSS T3 column (2.1 x 150 mm, 1.8µm) with an Acquity UPLC HSS T3 VanGuard Pre-Column (2.1 x 5 mm, 1.8µm) was utilized. The mobile-phase solvents consisted of solvent A = 10 mM ammonium formate, 5 μM deactivator additive (p/n 5191-3940, Agilent Technologies) in water/acetonitrile/isopropanol (5:3:2) and solvent B = 10 mM ammonium formate, 5 μM deactivator additive (p/n 5191-3940, Agilent Technologies) in water/acetonitrile/isopropanol (5:3:2). The column compartment temperature was maintained at 60° C and lipids were eluted using a linear gradient at a flow rate of 400 µL/ minute as follows: 0 minute 15% B, 2.5 minutes 50% B, 2.6 minutes 57% B, 9 minutes 70% B, 9.1 minutes 93% B, 11 minutes 96% B, 11.1 minutes 100% B, 12 minutes 100%B, 12.2 – 16 minutes 15% B. Data was acquired in positive and negative ion modes. The LC/MS data were then processed and analyzed using XCMS^35^, Compound Discoverer (version 3.3), and Skyline^34^. Statistical comparisons (unpaired t-tests, Benjamini and Hochberg false discovery rate <0.1), polar pathway enrichment with a hit rate threshold of ≥ 3 in MetaboAnalyst^36^, and lipid pathway analysis in BioPAN^37^. All samples collected for the metabolomics cohort were included in the analyses, with no exclusions.

### Sample size

This study was designed to define a hypothermic state induced by selective activation of a genetically defined neuronal population and to examine its association with ischemic injury, functional outcomes, and metabolic remodeling rather than to serve as a confirmatory preclinical efficacy study.

For <u>Experiment 1,</u> a total of N = 22 mice were initially used across experimental groups. After application of pre-specified physiological validity criteria, three mice were excluded due to failure to achieve adequate reperfusion. Six additional mice died from stroke severity prior to 24 hours and were therefore not available for serum and liver tissue collection; analyzed sample sizes thus vary accordingly. The remaining 13 mice (4 males, 9 females) were included in the final analyses.

For <u>Experiment 2</u>, a total of N = 53 mice were initially used across experimental groups. After application of pre-specified physiological validity criteria, six mice were excluded for failure to achieve hypothermia and four were excluded for failure to meet perfusion criteria. Additional mice died from stroke severity during the post-ischemic period; although these deaths were not applied as an exclusion criterion, mortality was itself treated as an outcome measure and compared across experimental groups. Analyzed sample sizes thus vary across other outcome measures depending on availability of downstream assessments. The remaining 43 mice (20 males, 23 females) were included in the final analyses.

For Experiment 3, a total of 17 mice (8 males, 9 females) were included in the metabolomics cohort with no animals excluded from the cohort.

## Statistics

Data are presented as mean ± 95% confidence interval (CI) or median with 95% CI, as appropriate. Normality was formally assessed for all datasets using the Shapiro–Wilk test prior to statistical analysis. Core body temperature reductions were analyzed, based on normality assumptions, using a one-way analysis of variance (ANOVA) or Kruskal–Wallis test to assess differences among the three experimental groups, followed by Tukey’s or Dunn’s multiple comparisons test for post hoc pairwise comparisons. For behavioral outcomes (rotarod, pole, and corner tests), stroke size and body weight loss, which did not meet normality assumptions, data were analyzed using the Kruskal–Wallis test, with Dunn’s multiple comparisons test for post hoc pairwise comparisons.

Post hoc exploratory sensitivity analyses were performed to assess potential sex-dependent treatment effects. For the 12-h temperature-change endpoint, infarct volume, and percent body-weight change, linear models incorporating sex, treatment, and the sex × treatment interaction were fitted and evaluated using Type III ANOVA. Longitudinal behavioral outcomes (Rotarod, Pole, and Corner tests) were analyzed using linear mixed-effects models with sex, treatment, and time as fixed effects, including all two- and three-way interactions, and mouse as a random intercept to account for repeated measurements within individual animals.

Longitudinal body-temperature data were similarly analyzed using linear mixed-effects models incorporating sex, treatment, time, and their interactions, with mouse included as a random intercept. Post hoc sex comparisons were estimated using estimated marginal means.

Mortality attributable to stroke severity was treated as an outcome measure and compared between experimental groups using Fisher’s Exact test, given small sample sizes. Mortality was not used as an exclusion criterion. Correlations between the extent of hypothermia (hypothermic index) and infarct size were evaluated at multiple time points using linear regression, reporting the coefficient of determination (r²) and associated p-values. All analyses and data visualization were performed using Prism (GraphPad Software, San Diego, CA). Significance was set at *P* < 0.05 unless otherwise specified.

For targeted metabolomic analyses of liver and serum, standardized effect sizes (Cohen’s d) and 95% confidence intervals were calculated for each metabolite comparing control and hypothermic-POA^KOR+^conditions. Confidence intervals were estimated using 10,000 bootstrap iterations with bias-corrected resampling.

Metabolomic data were analyzed using unpaired t-tests for exploratory screening between groups. A false discovery rate (FDR) correction was applied using the Benjamini–Hochberg method (threshold < 0.1) to identify statistically robust differences. Heatmaps were used to visualize global patterns of metabolite and lipid abundance, highlighting the top species showing the greatest variation across conditions. Pathway enrichment analyses were conducted to interpret the biological relevance of the identified metabolic and lipid changes. Pathway enrichment for polar metabolites was conducted using MetaboAnalyst^36^, with a hit rate threshold >3, and lipid pathway analysis was performed in BioPAN^37^ with a significance cutoff of |Z-score| > 0.95.

## Results

### Activation of POA^KOR^⁺ neurons induces sustained hypothermia with coordinated metabolic reprogramming after ischemic injury

We examined whether selective activation of POA^KOR+^ neurons induces sustained hypothermia with coordinated metabolic reprogramming in the setting of transient cerebral ischemia. Prior work in non-ischemic mice has demonstrated that POA^KOR^⁺ neuron activation induces coordinated hypothermic and hypometabolic responses^24^. Here, we used chemogenetic activation of POA^KOR+^ neurons after transient cerebral ischemia and quantified core body temperature trajectories and body metabolism for 24 hours (Fig. 1A).

**Figure 1.**
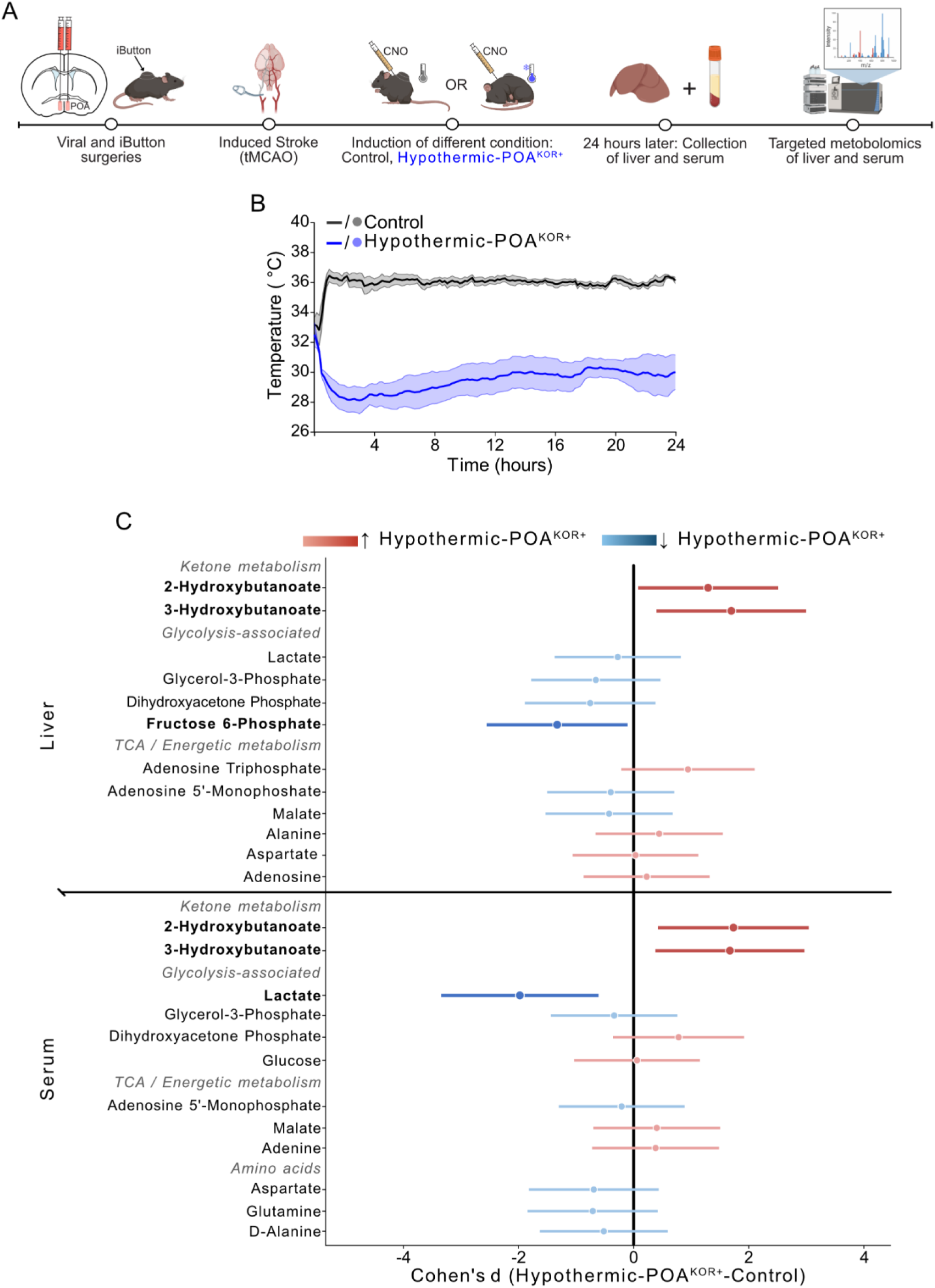
Selective activation of POA^KOR^⁺ neurons induces a sustained hypothermic state associated with coordinated systemic metabolic remodeling after ischemic injury. (A) Experimental design. KOR-Cre mice underwent stereotactic viral injection targeting POA^KOR+^ neurons and implantation of temperature-monitoring devices prior to transient middle cerebral artery occlusion (tMCAO). Following ischemia, mice received clozapine-N-oxide (CNO) to induce chemogenetic activation of POA^KOR+^ neurons under conditions permitting hypothermia (“POA^KOR+^”) or control conditions. Twenty-four hours later, serum and liver tissue were collected for targeted metabolomic analysis. (B) Core body temperature following POA^KOR+^-activation after tMCAO. Hypothermic-POA^KOR+^ mice exhibited a rapid and sustained reduction in core body temperature relative to controls throughout the 24-hour recording period. Shaded regions indicate mean ± SEM.

Core body temperature measurements revealed a sustained and selective hypothermic state (Fig. 1B). At 12 hours post-activation, mean absolute core body temperature was 28.61 °C (95% CI [26.71, 30.51] °C) compared to 35.93 °C (95% CI [35.17, 36.68] °C) in controls (one-way ANOVA, F(2,40) = 16.54, P < 0.0001; Tukey’s post hoc comparison, P < 0.0001). These differences were sustained across the full recording period, with significant group effects observed at 24 hours (one-way ANOVA, P = 1.19 × 10⁻⁴). A post hoc analysis identified a greater temperature reduction in males than females (sex × treatment interaction: F(1,9) = 9.60, P = 0.0127), although this analysis included only two males per treatment group and the interaction was not replicated in the larger Experiment 2 cohort.

Having established that POA^KOR+^ activation induces a sustained hypothermic state, we next examined whether this state was associated with coordinated systemic metabolic remodeling. Targeted metabolomic analysis of liver and serum focused on pathways related to glycolysis, the TCA cycle, ketone body metabolism, and amino acid metabolism. Figure 1C summarizes the effect sizes and confidence intervals for all measured metabolites. Across both liver and serum, glycolysis-associated metabolites generally shifted downward relative to controls, including lactate, glycerol-3-phosphate, fructose-6-phosphate, and dihydroxyacetone phosphate. Serum lactate demonstrated the largest negative effect size, consistent with reduced systemic glycolytic output in the hypothermic-POA^KOR+^ condition. This pattern extended to liver fructose-6-phosphate, an upstream glycolytic intermediate, supporting coordinated downregulation of glycolysis-associated metabolic pathways. In parallel, ketone body metabolites demonstrated some of the largest positive effect sizes across the dataset, with robust elevations in both serum and liver β-hydroxybutyrate species (2-hydroxybutyrate and 3-hydroxybutyrate). In contrast, metabolites associated with cellular energetic status and mitochondrial metabolism, including ATP, AMP, malate, and aspartate, generally showed comparatively smaller effect sizes centered closer to zero. Collectively, these findings define a coordinated hypothermic metabolic state characterized by reduced glycolysis-associated metabolites, increased ketone-associated metabolism, and relative preservation of central energetic pathways, with features resembling the coordinated fuel-switching observed during natural torpor and hibernation^14,17^.

Forest plot summarizing standardized effect sizes (Cohen’s d) and 95% confidence intervals for targeted metabolites measured in liver and serum comparing hypothermic-POA^KOR+^ and control conditions. Positive values indicate increased relative abundance in the hypothermic state, whereas negative values indicate reduced relative abundance. Dark colors indicate metabolites with confidence intervals not overlapping zero, whereas lighter colors indicate confidence intervals overlapping zero. Confidence intervals were estimated using 10,000 bootstrap iterations with bias-corrected resampling. Metabolites are grouped by biologic pathway category. Abbreviations: AMP, adenosine monophosphate; ATP, adenosine triphosphate; DHAP, dihydroxyacetone phosphate; F6P, fructose-6-phosphate; G3P, glycerol-3-phosphate; 2-HB, 2-hydroxybutyrate; 3-HB, 3-hydroxybutyrate.

### A POA^KOR^⁺ induced hypothermic state is associated with reduced ischemic injury and improved motor outcomes

We then asked if this hypothermic state with metabolic reprogramming was associated with cerebroprotection and whether this effect was temperature dependent. We therefore compared three experimental groups: (1) control mice injected with a control virus, housed at a thermoneutral temperature (30 °C) after tMCAO, (2) POA^KOR+^-activated mice housed at 30 °C after tMCAO, termed “hypothermic-POA^KOR+^”; and (3) POA^KOR+^ activated mice housed at an ambient temperature of 36 °C after tMCAO to minimize the development of hypothermia, termed “normothermic-POA^KOR+^”. By manipulating ambient temperature during neuronal activation we separate the effects of POA^KOR^⁺ activation from those attributable to reduced body temperature following ischemic injury.

Mice underwent 30-minute tMCAO prior to chemogenetic activation, and core body temperature was monitored for 72 hours prior to behavioral testing and histology (Fig. 2A). Core body temperature measurements revealed a sustained and selective hypothermic response in the hypothermic-POA^KOR+^ group housed at 30 °C. Within the first 12 hours post-activation, this group exhibited a mean temperature reduction of 3.35 °C (95% CI [2.50, 4.19] °C), compared to 0.55 °C (95% CI [0.05, 1.06] °C) in the control group and 1.73 °C (95% CI [0.53, 2.92] °C) in the normothermic-POA^KOR+^-activated group. Temperature change differed significantly across groups (*P* < 0.0001); post hoc testing indicated greater hypothermia in hypothermic-POA^KOR^⁺ mice relative to both control and normothermic-POA^KOR+^ groups (Fig. 2B). Temperatures in the hypothermic-POA^KOR+^ group gradually returned toward baseline after 24 hours, whereas the other two groups maintained relatively stable temperatures throughout the monitoring period. In contrast to Experiment 1, there was no sex × treatment interaction for the 0–12-hour temperature response in this larger cohort (F(2,37) = 0.22, P = 0.804).

**Figure 2.**
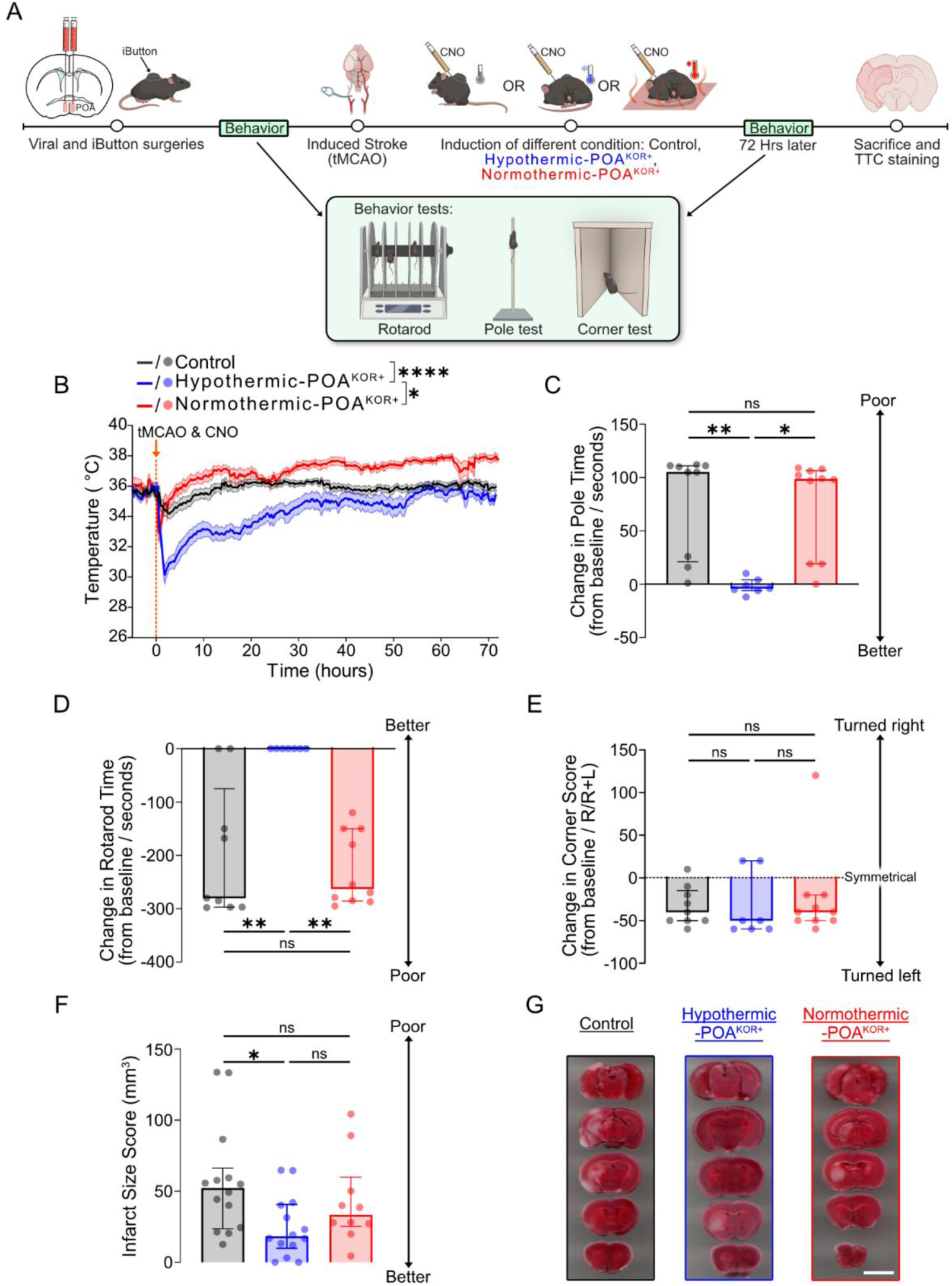
Activation of POA^KOR+^ neurons induces a sustained hypothermic state and is associated with improved outcomes following stroke. (A) Experimental timeline. KOR-Cre mice underwent viral injection followed by iButton implantation for temperature monitoring, baseline behavioral testing, and a 30-minute transient middle cerebral artery occlusion (tMCAO), with clozapine-N-oxide (CNO) injection to induce a hypothermic state in control (N = 17), hypothermic-POA^KOR^⁺ (N = 16), or normothermic-POA^KOR^⁺ (N = 10) conditions. Animals were reassessed at 72 hours using behavioral testing and histology. (B) Core body temperature trajectories over 72 hours following tMCAO. Group differences in temperature reduction (mean reduction over the first 12 hours post-activation) were assessed by one-way ANOVA F(2,40) = 16.54, P < 0.0001 with Tukey’s post hoc tests (*P<0.05, ***P<0.0001). (C) Change in pole test performance, calculated as the difference between baseline and 72 hours post-stroke. Group differences were assessed by Kruskal–Wallis test with Dunn’s post hoc comparisons (*P < 0.05, **P < 0.01). (D) Change in rotarod performance, calculated as the difference between baseline and 72 hours post-stroke. Group differences were assessed by Kruskal–Wallis test with Dunn’s post hoc comparisons (**P < 0.01). (E) Corner test asymmetry, expressed as the percentage of right turns relative to total turns (right + left). Group differences were assessed by Kruskal–Wallis test. (F) Quantification of infarct size using direct measurement, comparing the hypothermic-POA^KOR+^, control groups and normothermic-POA^KOR+^ groups. (Control N = 14; Hypothermic-POA^KOR+^ N = 14; Normothermic-POA^KOR+^ N = 10). One-way ANOVA, F(2,35) = 3.75, P < 0.05 by Tukey’s multiple comparison test. (G) Representative TTC (2,3,5-triphenyltetrazolium chloride) stained coronal sections from 3 individual mice, showing unstained infarct regions. Note: behavioral analyses (C–E) were performed on subsets of animals: Hypothermic-POA^KOR+^ (N = 7), Normothermic-POA^KOR+^ (N = 10), and Control (N = 9).

Under conditions permitting POA^KOR^⁺–induced hypothermia, animals exhibited reduced post-stroke motor impairment, whereas normothermic POA^KOR^⁺ mice exhibited deficits comparable to controls. In the pole test, hypothermic-POA^KOR+^ mice showed minimal change from baseline (median change = –4.0 sec, 95% CI [–12, 10]), in contrast to control (median change = 105.0 sec, 95% CI [16, 111] and to normothermic-POA^KOR+^ groups (median change = 98.5 sec, 95% CI [19, 108]; *P* = 0.0010 Fig. 2C). A similar pattern emerged in the rotarod test, where hypothermic-POA^KOR+^ mice exhibited minimal performance decline relative to baseline (median change = 0.0 sec, 95% CI [0, 0]), whereas control mice declined significantly (median change = –280.0 sec, 95% CI [0, –297]) and normothermic-POA^KOR+^ mice also performed poorly (median change = –262.5 sec, 95% CI [–150, –287]; *P* = 0.0021; Fig. 2D). All three groups demonstrated persistent post-stroke asymmetry on the corner test, with no significant differences among groups (hypothermic-POA^KOR+^ median = –50.0%, 95% CI [–60, 20]; control median = –40.0%, 95% CI [–50, –10]; normothermic-POAKOR+ median = –40.0%, 95% CI [–50, –20]; *P* = 0.5872; Fig. 2E).

Infarct size was markedly reduced in the hypothermic-POA^KOR+^ group (mean = 24.82 mm³, 95% CI [12.52, 37.13]) compared to controls (mean = 56.69 mm³, 95% CI [34.78, 78.61]), and was numerically smaller than the normothermic-POA^KOR+^ group (mean = 42.91 mm³, 95% CI [20.70, 65.13]; *P* = 0.0334; Fig. 2F, G). Post hoc sensitivity analyses showed no evidence that sex modified the treatment effect on infarct volume (sex × treatment, P = 0.745) or behavioral outcomes (sex × treatment × time: Rotarod, P = 0.341; Pole, P = 0.845; Corner, P = 0.178). Mortality rates at 72 hours were 17.65% in the control group (3/17), 18.75% in the hypothermic-POA^KOR+^ group (3/16), and 0% in the normothermic-POA^KOR+^ group (0/10); *P* = 0.4702, indicating that differences in infarct size and functional performance are unlikely to be explained by differential survivorship. Body weight loss was comparable across groups: hypothermic-POA^KOR+^ mice lost a median of 18.30% (95% CI [–21.40%, –15.90%]), control mice lost 12.40% (95% CI [–28.30%, –5.30%]), and normothermic-POA^KOR+^ mice lost 17.55% (95% CI [–23.70%, –4.40%]) (*P* = 0.8991). There was also no sex × treatment interaction for body weight change (F(2,31) = 0.025, P = 0.976).

We next assessed whether the magnitude and duration of hypothermia were associated with infarct size, using the hypothermic index, defined as the area under the temperature curve relative to baseline. A significant correlation was observed between the hypothermic index and stroke size at 6 hours (r^2^ = 0.18, *P* = 0.008), 12 hours (r^2^ = 0.22, *P* = 0.003; Fig. 3A), 24 hours (r^2^ = 0.23, *P* = 0.002) and 48 hours (r^2^ = 0.143, *P* = 0.019) post-stroke, but not 72 hours (Fig. 3B). Thus, across animals, greater depth and duration of hypothermia were associated with smaller infarct volumes during the acute post-stroke period. These findings indicate that the POA^KOR^⁺ induced hypothermic state is associated with smaller infarct volumes during the acute post-stroke period, consistent with a therapeutic window concentrated within the first 48 hours after ischemic injury.

**Figure 3.**
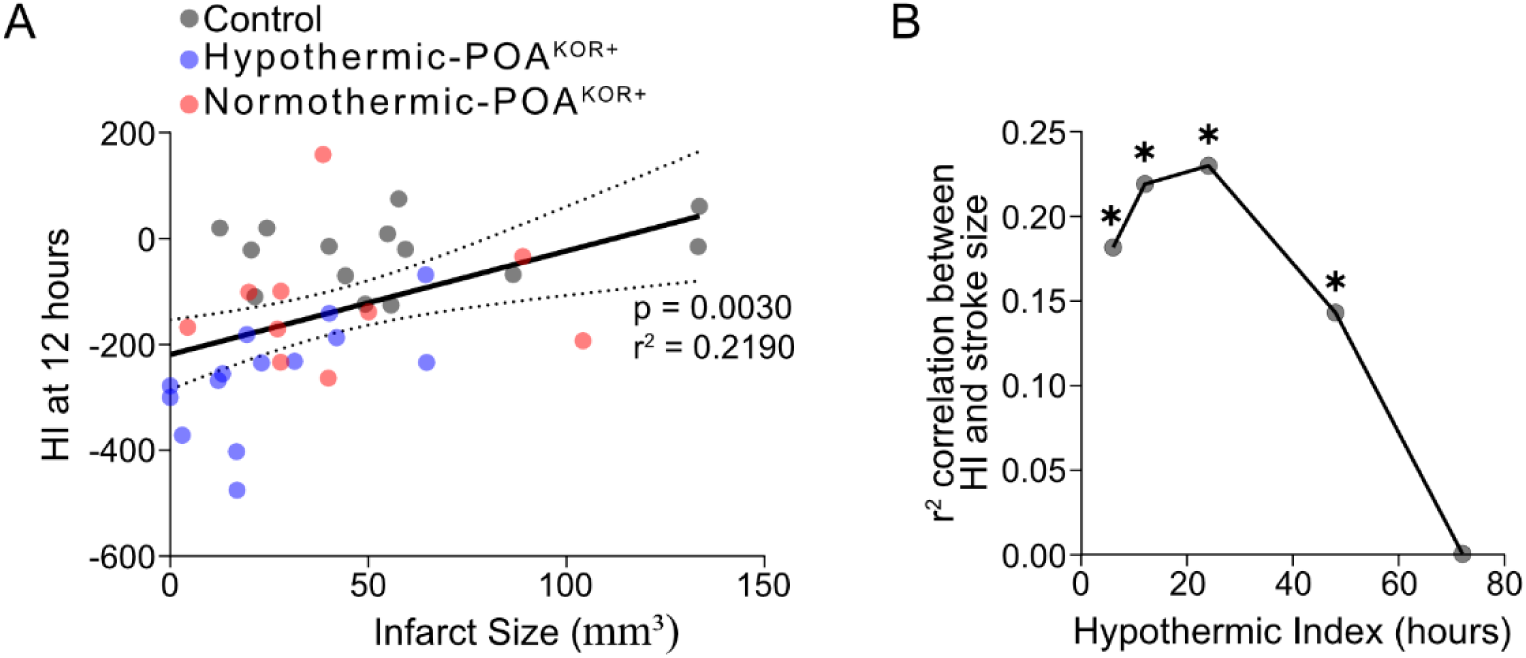
Magnitude of the hypothermic state is associated with acute stroke size. (A) The hypothermic index (HI), defined as the area under the core body temperature curve relative to baseline, was calculated to capture the depth and duration of hypothermia during the first 12 hours following stroke and plotted against direct infarct volume. (B) Associations between HI calculated over multiple post-stroke intervals (6, 12, 24, 48, and 72 hours) and infarct size are shown. Coefficients of determination (r²) indicate the strength of association at each time point. *P < 0.05.

### Metabolic features of the POA^KOR^⁺ induced hypothermic state

To define metabolic features associated with a cerebroprotective hypothermic state, we performed untargeted metabolomic profiling of brain tissue following POA neuronal activation under hypothermic versus normothermic conditions. To isolate early metabolic features, whole-brain tissue was collected 30 minutes after POA^KOR^⁺ activation, prior to ischemic injury or inflammatory sequelae. We compared three groups: (1) hypothermic-POA^KOR+^ mice, housed at room temperature (∼22 °C), that developed a hypothermic state following activation; (2) normothermic-POA^KOR+^ mice, housed at an elevated ambient temperature (36 °C), in which hypothermia was prevented; and (3) control mice injected with control virus and housed at room temperature (Fig. 4A). Core body temperature measurements confirmed selective hypothermia in the hypothermic-POA^KOR^⁺ group, whereas normothermic-POA^KOR^⁺ and control animals maintained stable temperatures (Fig. 4B). This design enabled direct comparison of metabolic features associated with hypothermia versus POA^KOR^⁺ activation in the absence of temperature reduction.

**Figure 4.**
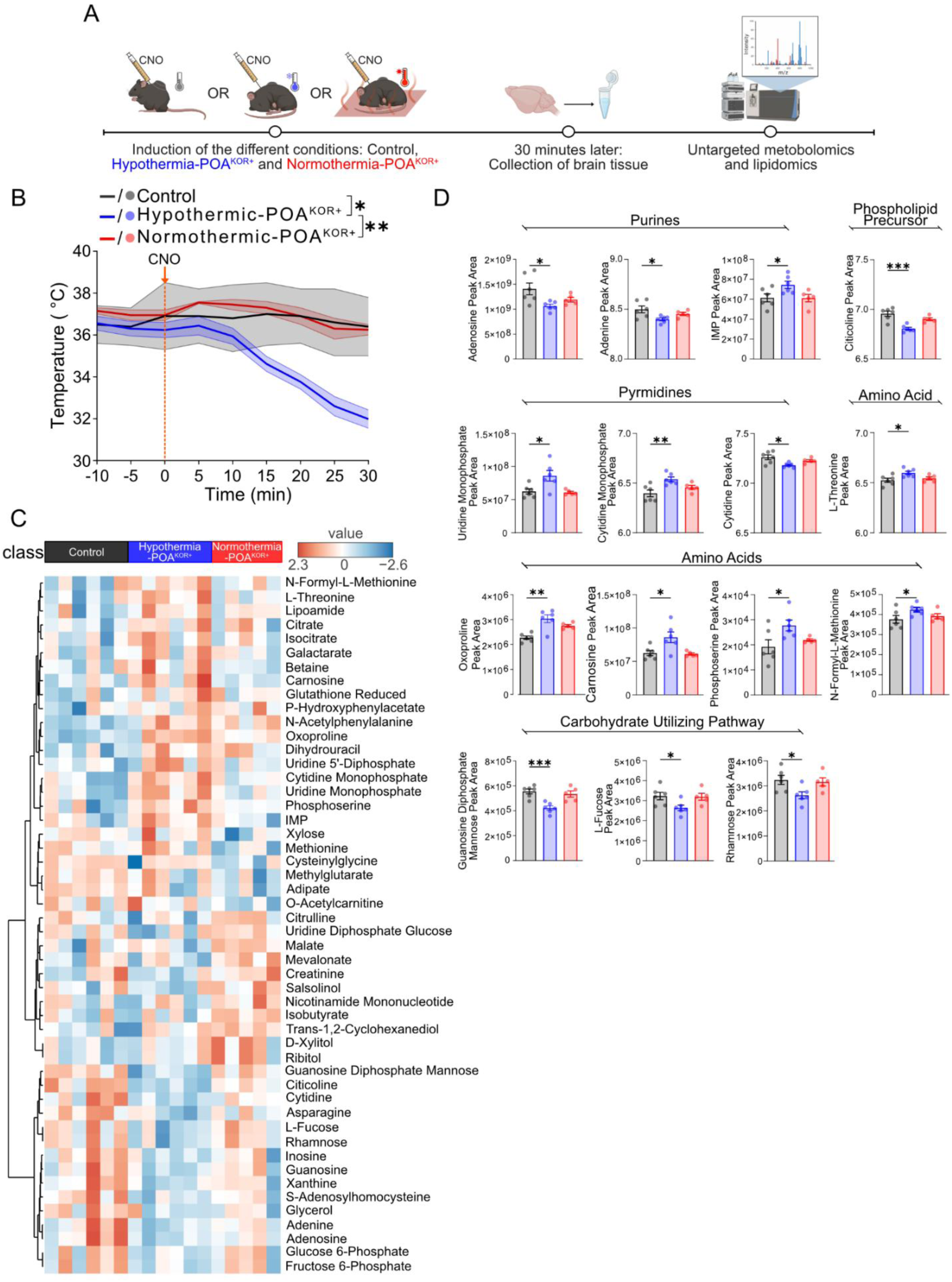
Early metabolomic changes associated with a hypothermic state following POA^KOR+^ activation. (A) Experimental design for untargeted metabolomics and lipidomics. Three groups were analyzed: hypothermic-POA^KOR^⁺ mice housed at room temperature (∼22 °C; N = 6), normothermic-POA^KOR^⁺ mice housed at 36 °C (N = 5), and controls housed at room temperature (N = 6). (B) Core body temperature traces during the 30 minutes following CNO injection for each experimental group. *P < 0.05, **P < 0.01 by Kruskal–Wallis test with Dunn’s post hoc comparisons. (C) Heatmap of metabolite abundance across conditions for the top 50 polar metabolites showing the greatest variance across conditions. Metabolite intensities were log10-transformed and Z-score normalized prior to visualization. Red indicates higher relative abundance and blue indicates lower relative abundance for each metabolite. (D) Fifteen metabolites identified using an uncorrected screening threshold (*P < 0.1, **P < 0.05, ***P < 0.01) in the hypothermic-POA^KOR^⁺ group relative to both control and normothermic-POA^KOR^⁺ groups. Abbreviation: IMP (inosine monophosphate).

We first examined global metabolic patterns using heatmaps of the top 50 polar metabolites that best distinguished the experimental groups (Fig. 4C). In control animals, metabolic profiles were relatively uniform, reflecting a stable baseline metabolic state. In contrast, hypothermic-POA^KOR^⁺ mice exhibited a distinct and coordinated shift across multiple metabolite classes, defining a metabolic profile associated with the hypothermic state following neuronal activation. Normothermic-POA^KOR^⁺ mice displayed profiles that either resembled controls or shifted in a pattern distinct from hypothermic-POA^KOR^⁺ mice, indicating that preventing hypothermia alters the metabolic state associated with POA^KOR^⁺ activation.

We next performed statistical comparisons to identify specific polar metabolites associated with the hypothermic state under POA^KOR^⁺ activation. We compared the hypothermic-POA^KOR^⁺ group to both normothermic-POA^KOR^⁺ and control groups, using an initial uncorrected screening threshold (P < 0.1) to avoid prematurely excluding potentially meaningful changes.

This exploratory analysis identified 15 polar metabolites selectively altered in the hypothermic-POA^KOR^⁺ group, but not in normothermic or control animals. Pathway enrichment analysis of these metabolites revealed predominant involvement of purine and pyrimidine metabolism, phospholipid biosynthesis, and carbohydrate-utilizing pathways (Fig. 4D). These pathways encompass nucleotide salvage, membrane phospholipid turnover, and biosynthetic adaptations to cellular stress, that have been associated with resilience under ischemic conditions ^35–39^. To identify the most robust changes, we then applied FDR correction (threshold < 0.1) across all measured metabolites. Three metabolites—citicoline, guanosine diphosphate mannose, and oxoproline—remained significant after correction. While the remaining twelve metabolites did not meet FDR significance, they showed directionally consistent changes and are included in Fig. 4D for reference. These analyses define a coordinated set of metabolic features associated with the neural pathway induced hypothermic state.

We next performed untargeted lipidomic profiling on whole-brain extracts from the same experimental groups. We first generated a heatmap of the top 50 lipids showing the greatest variation across conditions (Fig. 5A). Consistent with the polar metabolite analysis, hypothermic-POA^KOR^⁺ mice exhibited widespread, coordinated lipid changes, while normothermic-POA^KOR^⁺ and control samples showed more stable profiles.

**Figure 5.**
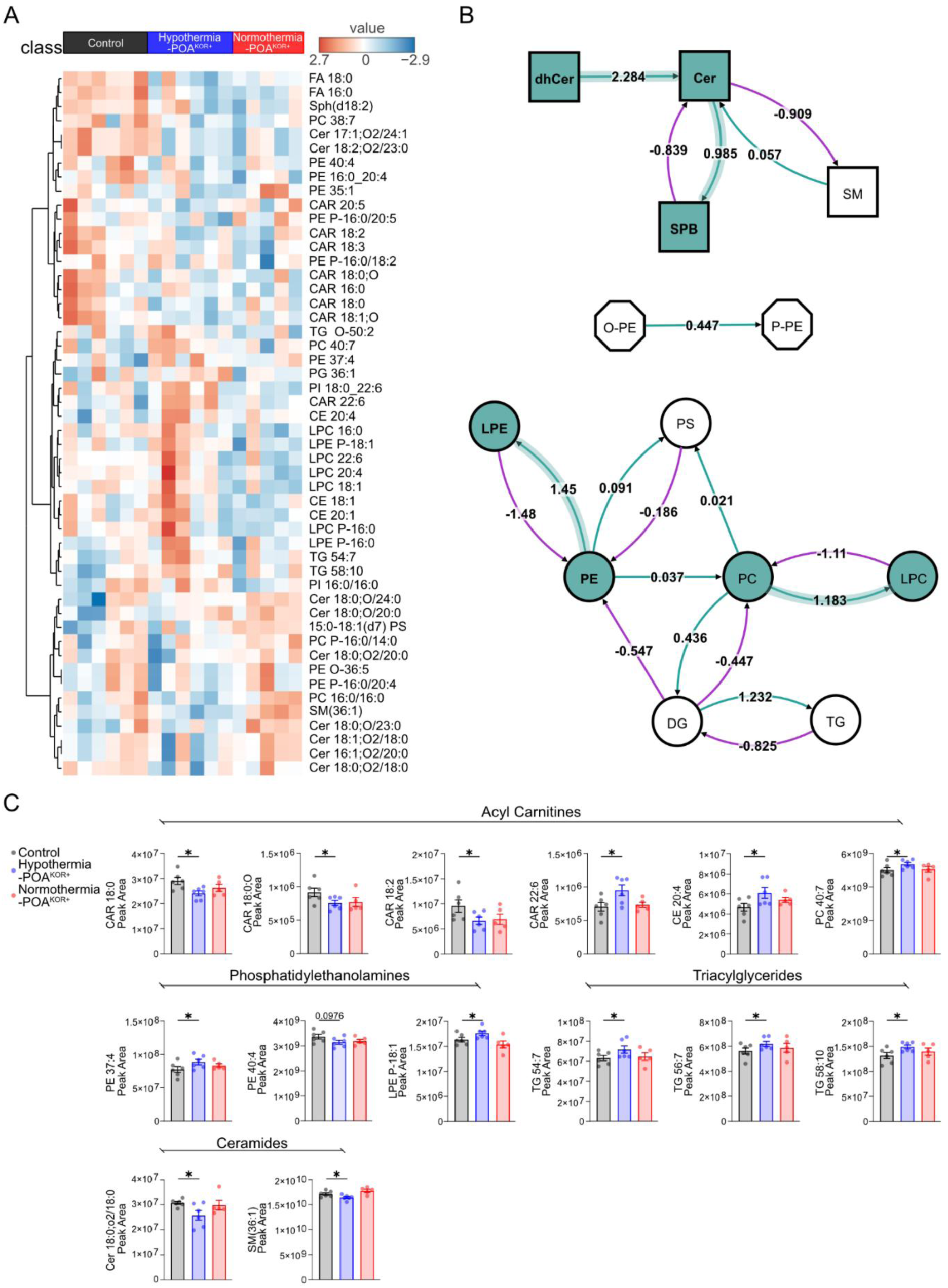
Lipidomic profiling associated with the hypothermic state following POA^KOR+^ activation. (A) Heatmap of lipid abundance across conditions for the top 50 lipids showing the greatest variance across groups (hypothermic, N = 6; normothermic, N = 5; control, N = 6). (B) Pathway analysis of lipid metabolism. Nodes represent lipid classes. Edges indicate transformations between lipid species, with numbers representing relative Z-score changes in inferred lipid conversion. Cyan nodes and arrows indicate pathways showing increased inferred conversion in the hypothermic-POA^KOR^⁺ group relative to control, but not normothermic-POA^KOR^⁺ conditions, using a threshold of |Z-score| > 0.95. White nodes and unshaded arrows indicate pathways with no detectable change in conversion. (C) Fourteen lipid species identified using an uncorrected screening threshold (uncorrected *P < 0.1) in the hypothermic-POA^KOR^⁺ condition relative to control, but not normothermic-POA^KOR^⁺ conditions. Lipid class abbreviations: dhCer (Dihydroceramide), Cer (Ceramide), SBP (Sphingosine), SM (Sphingomyelin), PE (Phosphoethanolamine), PS (Phosphoserine), PC (Phosphocholine), DG (Diacylglycerol), TG (Triacylglycerol), FA (Fatty Acid), Sph (Sphinganine), CAR (Carnitine), PE P (Plasmalogen Phosphoethanolamine), PG (Phosphoglycerol), PI (Phosphatidylinositol), CE (Cholesteryl Ester), LPC (Lysophosphatidylcholine), LPE (Lysophosphatidylethanolamine), O-PE (Ether-linked Phosphoethanolamine), P-PE (Plasmenyl-Phosphoethanolamine).

We then performed pathway-level analysis to identify lipid pathways associated with the hypothermic state following POA^KOR+^ activation (Fig. 5B). This analysis compared hypothermic-POA^KOR^⁺ mice to controls and evaluated whether identified pathways remained unchanged when hypothermia was prevented under normothermic conditions. This comparison highlighted lipid pathways altered in the hypothermic-POA^KOR^⁺ group, including sphingolipid processing (e.g., conversion of dihydroceramides to ceramides, increased sphingoid base production), phosphatidylcholine breakdown, and lysophospholipid remodeling. These pathways encompass processes related to membrane remodeling, stress response, and cellular energetics, and that have been linked to resilience mechanisms during ischemic stress^43–49^. Following pathway analysis, we screened individual lipid species for differences between groups using an initial uncorrected P-value threshold of < 0.1. Although no individual lipids met significance after FDR correction, multiple lipid classes, including acylcarnitines, cholesteryl esters, ceramides, sphingomyelins, phosphatidylethanolamines, and triglycerides, exhibited directionally consistent changes selectively enriched in the hypothermic condition (Fig. 5C). Together, these lipidomic findings define temperature-dependent alterations in membrane-associated lipid pathways during the hypothermic state.

## Discussion

Activation of POA^KOR^⁺ neurons produced a sustained hypothermic state with metabolic reprogramming following transient cerebral ischemia and was associated with reduced infarct size and improved acute motor outcomes. When hypothermia was prevented despite neuronal activation, these protective effects were attenuated, indicating that temperature reduction is necessary for cerebroprotection. Concurrent untargeted metabolomic profiling revealed coordinated, temperature-dependent remodeling of nucleotide, phospholipid, and sphingolipid pathways, demonstrating that the induced hypothermic state involves structured metabolic reorganization.

These results align with prior studies demonstrating that stimulation of neuronal circuits in the POA are sufficient to induce hypothermia and reduce ischemic brain injury^21,22^. Building on this body of work, the present study refines existing models by identifying a genetically specified POA^KOR^⁺ neuronal subpopulation^20^ whose activation induces a sustained hypothermic state and coordinated metabolic reprogramming in the context of ischemic injury. To characterize the metabolic features of the induced state, we performed untargeted metabolomic profiling shortly after POA^KOR^⁺ activation to capture early biochemical changes. Under hypothermic conditions, POA^KOR^⁺ activation produced coordinated changes across metabolic pathways involved in nucleotide metabolism, phospholipid remodeling, and lipid processing. These pathway-level alterations were largely attenuated or directionally distinct when hypothermia was prevented.

The metabolic pathways altered in the induced hypothermic state have been linked to cellular resilience under ischemic stress. Alterations in purine and pyrimidine metabolism are consistent with prior work suggesting that enhanced nucleotide salvage is associated with preservation of energy balance and macromolecular integrity during ischemia, including maintenance of ATP availability and nucleotide pools^40,41^. Changes in phospholipid-related metabolites, including citicoline (a compound previously associated with cerebroprotection in experimental and clinical stroke studies^50–54)^ as well as lysophosphatidylcholine species, are consistent with adaptive remodeling of membrane composition and signaling pathways implicated in neuronal survival and inflammatory resolution^55–57^. In parallel, the observed shifts in sphingolipid-related pathways are notable given evidence that ischemic injury is associated with accumulation of pro-apoptotic ceramides, whereas diversion toward alternative sphingolipid species is associated with stress adaptation and cell survival^17,48^.

Taken together, these findings support the broader concept that engaging thermoregulatory circuits establishes a hypothermic state with metabolic reprogramming conceptually distinct from externally imposed cooling. Conventional hypothermia strategies reduce body temperature by opposing homeostatic defenses, often eliciting counter-regulatory responses such as shivering and increased metabolic demand. In contrast, activation of POA^KOR^⁺ neurons engages intrinsic thermoregulatory programs, producing a coordinated hypothermic state associated with structured metabolic remodeling. This framing is consistent with prior work demonstrating that stimulation of central thermoregulatory neural populations, including within the POA, can be sufficient to induce hypothermia and reduce ischemic brain injury^21,22^, as well as emerging evidence that torpor-like states confer organism-level benefits beyond acute injury contexts^58^.

Several limitations of the present study should be acknowledged. Stroke outcomes were assessed under thermoneutral housing (30 °C), consistent with standard preclinical stroke protocols that minimize thermoregulatory stress, whereas metabolomic profiling was conducted at room temperature to allow physiological expression of the induced hypothermic state.

Importantly, mice in both paradigms began at comparable core body temperatures and exhibited similar magnitudes of POA^KOR+^-induced hypothermia. Nevertheless, the distinct ambient conditions constrain direct causal linkage between metabolic features and ischemic protection. In addition, metabolomic analyses were performed on postmortem brain tissue, and highly labile metabolites may be influenced by postmortem metabolism prior to tissue freezing. Accordingly, although absolute *in vivo* metabolite concentrations cannot be inferred, all experimental groups were collected and processed using identical procedures, supporting interpretation of the findings as relative differences between groups. Physiological parameters such as blood glucose, arterial and venous oxygen saturation, and circulating stress hormones were also not measured in the present study. Future studies incorporating additional physiological measures, including blood glucose, arterial and venous oxygen saturation, and circulating stress hormones, as well as metabolic profiling at multiple time points across the evolution of the hypothermic state, will provide a more comprehensive characterization of the systemic response and its temporal dynamics. Although both sexes were included, the study was not designed or powered to establish sex-specific effects. Exploratory analyses identified a potential sex difference in the hypothermic response that was not replicated in the larger cohort, while providing no evidence that sex modified the principal cerebroprotective outcomes. Isoflurane exposure may also modify the physiological context in which POA^KOR+^ activation occurs, including through effects on thermoregulatory circuitry and ischemic injury. Although all stroke groups received equivalent isoflurane exposure and POA^KOR+^ activation was initiated after discontinuation of anesthesia, the present design does not exclude an interaction between peri-anesthetic exposure and the magnitude or characteristics of the induced hypothermic state. Finally, the study focuses on acute injury and functional outcomes within a 72-hour window and does not address durability of protection or longer-term recovery. Thus, the present work does not establish specific causal metabolic mechanisms or long-term efficacy.

KOR-expressing neurons have been identified within the human POA, suggesting that analogous thermoregulatory architectures exist in the human brain^25,26^. Additionally, recent work has demonstrated that torpor-like hypothermic states can be induced in non-hibernating mammals using noninvasive neuromodulatory approaches, including focused ultrasound, indicating that engagement of central thermoregulatory circuits is technically feasible without genetic manipulation^59^. Together, these observations support the plausibility that conserved thermoregulatory circuits can be targeted to induce controlled hypothermic states.

This study establishes that selective activation of a genetically defined thermoregulatory circuit in the POA is sufficient to generate a sustained, hypothermic state associated with reduced acute ischemic injury. These findings emphasize engagement of thermoregulatory circuits as a framework for understanding hypothermia-associated brain protection at the level of biological state. The accompanying metabolic signatures provide a descriptive foundation for future studies aimed at testing how neural circuit-defined regulation of temperature and metabolism shapes tissue vulnerability and recovery trajectories during brain injury.

## Acknowledgments

The authors gratefully acknowledge Spencer Blackwood and Hee Ra Jung for their assistance with the experiments. All scientific content was written, verified, and approved by the authors. This work was supported by the National Institutes of Health (NIH), National Institute of Neurological Disorders and Stroke (NINDS) under the following grants: R01 NS133365-01, R37 NS110699, and R01 NS120481.

## Author contributions

Conceptualization: ECL, AJN Methodology: CY, AK, ERG, JL, HES, GJP Visualization: KMI, AK

Animal Surgery: ERG, TAA, JL, AK, HES Funding acquisition: ECL, AJN, JML Data collection: AK, JKC, JL, ERG Statistical Analysis: AK, FZ

Supervision: ECL, AJN, LPS, JML Writing –original draft: AK, ECL

Writing – review & editing: AK, ECL, AJN, LPS, JML, JKC, TAA

## Data and materials availability

Raw metabolomics data have been deposited in the Metabolomics Workbench (https://www.metabolomicsworkbench.org) and will be released upon publication. The accession number will be added once it is available. All other data supporting the findings of this study are included in the article and its supplementary information or are available from the corresponding author upon reasonable request.

