## Supplemental Figures S1 and S2 for "Targeted preoptic neuronal activation induces a hypothermic state with metabolic reprogramming and acute cerebroprotection"

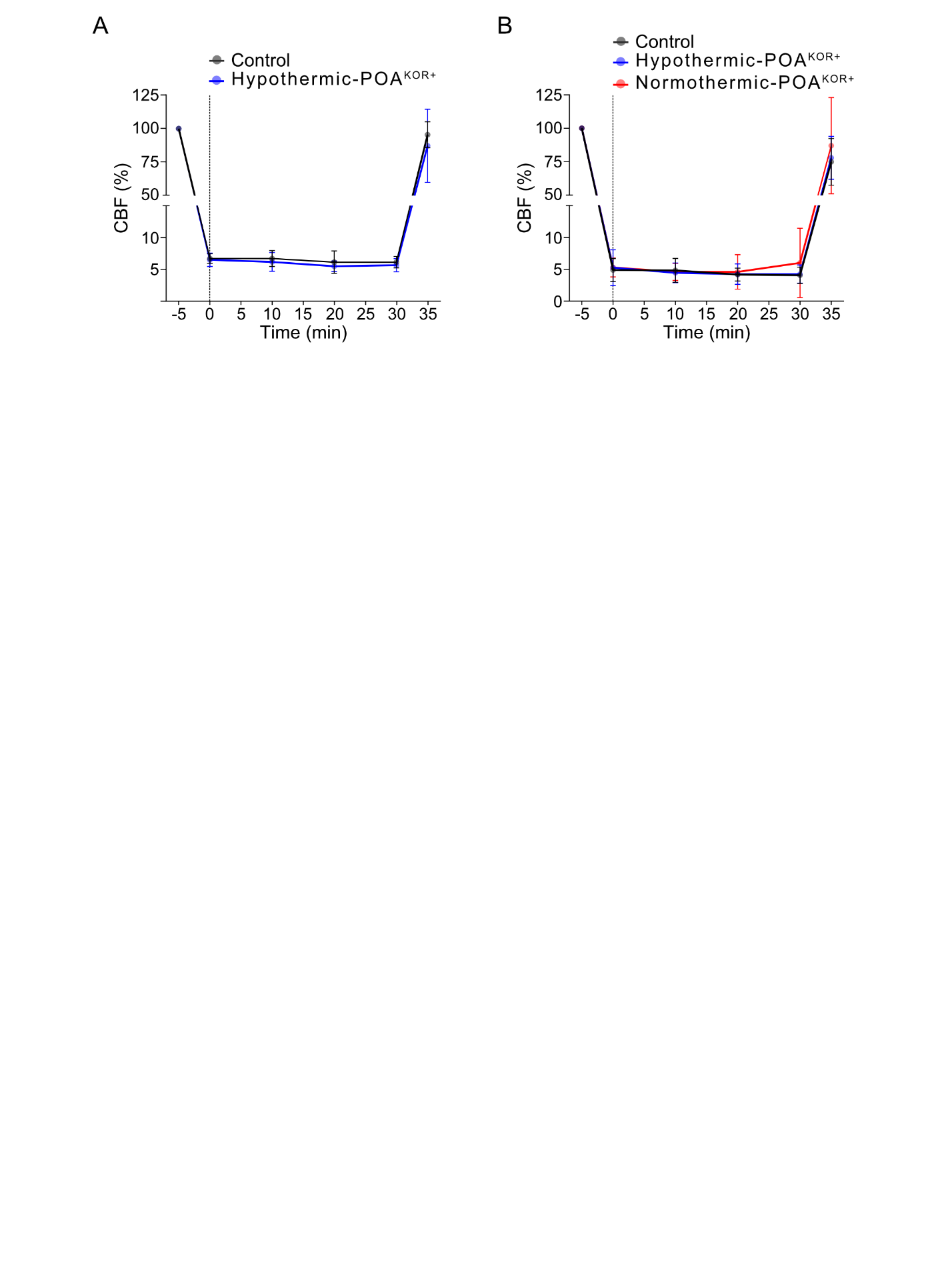

**Supplementary Figure S1.** **Cerebral blood flow monitoring during transient middle cerebral artery occlusion (tMCAO).**
**(A) Experiment 1.** Relative cerebral blood flow (CBF, % baseline) measured by laser Doppler flowmetry before occlusion, during 30 min of tMCAO, and following reperfusion in Control and Hypothermic- POA^KOR+^mice. **(B)** **Experiment 2.** Relative CBF profiles in Control, Hypothermic- POA^KOR+^and Normothermic- POA^KOR+^mice. Data is presented as mean ± SD. The vertical dashed line indicates the onset of MCA occlusion (time = 0 min).

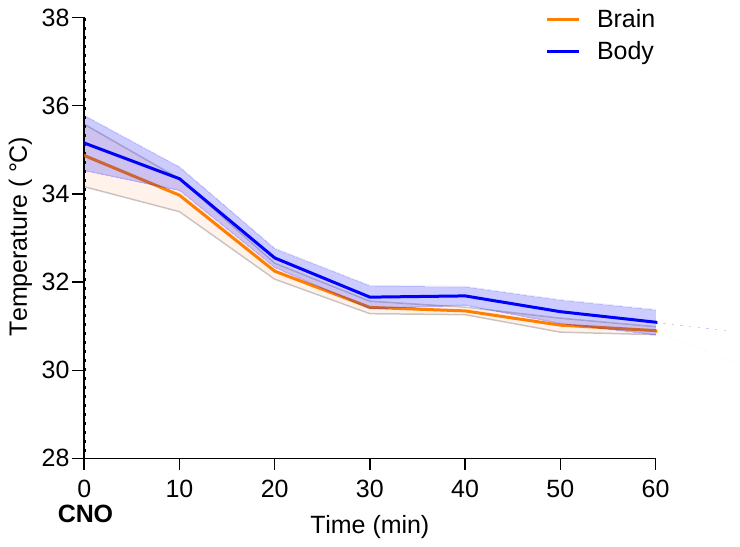

**Supplementary Figure S2. Brain and body temperature during POA^KOR^⁺-induced hypothermia (N = 4).** Simultaneous recordings of brain and subcutaneous body temperature following a single CNO administration show parallel temporal decreases. Data are presented as mean ± SEM.
